# *Ankhd1* enhances polycystic kidney disease development via promoting proliferation and fibrosis

**DOI:** 10.1101/2020.03.04.977017

**Authors:** Foteini Patera, Guillaume M Hautbergue, Patricia Wilson, Paul C Evans, Albert CM Ong, Maria Fragiadaki

## Abstract

Autosomal Dominant Polycystic Kidney Disease (ADPKD) is the most common genetic kidney disorder resulting in 10% of patients with renal failure. The molecular events responsible for the relentless growth of cysts are not defined. Thus, identification of novel drivers of ADPKD may lead to new therapies. Ankyrin Repeat and Single KH domain-1 (ANKHD1) controls cancer cell proliferation, yet its role in ADPKD is unexplored. Here, we present the first data that identify ANKHD1 as a driver of proliferative growth in cellular and mouse models of ADPKD. Using the first *Ankhd1*-deficient mice, we demonstrate that *Ankhd1* heterozygosity potently reduces cystic growth and fibrosis, in a genetically orthologous mouse model of ADPKD. We performed transcriptome-wide profiling of patient-derived ADPKD cells with and without ANKHD1 siRNA silencing, revealing a major role for ANKHD1 in the control of cell proliferation and matrix remodelling. We validated the role of ANKHD1 in enhancing proliferation in patient-derived cells. Mechanistically ANKHD1 promotes STAT5 signalling in ADPKD mice. Hence, ANKHD1 is a novel driver of ADPKD, and its inhibition may be of therapeutic benefit.

## INTRODUCTION

Autosomal Dominant Polycystic Kidney Disease (ADPKD) is the most common genetic cause of renal failure, characterised by hyperproliferation of renal tubular epithelial cells^1,2^. In addition, patients develop liver and pancreatic cysts and experience intracranial aneurysms at a significantly increased rate than the general population (10%). Over 12 million people are affected with ADPKD worldwide^3^. Hence ADPKD is a multi-organ disease with significant socio-economic impact that currently lacks a cure. A vasopressin receptor antagonist (tolvaptan) is the only approved medicine to slow down ADPKD progression, demonstrating proof-of-principle that the disease can be modified. Yet the downside of tolvaptan are the severe side effects (e.g. liver failure requiring liver transplantation and diabetes insipitus), highlighting the need new for new medicines. Discovery of new ways to treat ADPKD depends on a molecular level understanding of the pathobiology underlying ADPKD progression, defined by the relentless growth of cysts.

Molecularly, the initiation of ADPKD is due to mutations in the *PKD1* or *PKD2* genes ^4,5^ leading to striking tissue overgrowth, resembling cancer. The cause of the relentless cystic growth, once disease is established, is however unclear. Next generation sequencing approaches have recently provided strong evidence of the involvement of altered RNA metabolism in the pathogenesis of ADPKD^6,7^. Despite the significantly improved understanding achieved from these powerful studies, we still do not know the proteins responsible for regulating RNA metabolism in ADPKD. RNA binding proteins (RBPs) control all aspects in the life cycle of RNA, from synthesis to splicing and degradation^8^. An example of an RBP with a known role in ADPKD is bicaudal-c homolog 1 (Bicc1), which is the gene responsible for the jcpk and bpk spontaneous mouse models of polycystic kidney disease^9^. Bicc1 contains three classical KH RNA binding domains and a SAM RNA recognition domain, which together allow Bicc1 to recognize and bind its specific RNA targets^10^. Another RBP that has never been studied in ADPKD is Ankyrin Repeat and Single KH domain 1 (ANKHD1). ANKHD1 has a classic RNA binding domain (KH) and is involved in proliferative signalling in *Drosophila*, by enhancing for example JAK/STAT, HIPPO and EGF/MAP pathway activities^11–14^. Interestingly all of aforementioned signalling pathways are dysregulated and drive ADPKD development^15–18^. Moreover, ANKHD1 enhances renal cancer cell growth^19^. Therefore, it is hypothesized that ANKHD1 may play a role in disease processes in ADPKD, by controlling the way epithelial cells with Pkd1 mutations proliferate.

Ankyrin Repeat and single KH domain 1 (ANKHD1) is a protein with an evolutionarily conserved KH domain, which binds RNA to promote cell growth of renal cell carcinoma^20^, yet its role of ADPKD is unknown. The *Drosophila* homologue of ANKHD1, *MASK*, enhances JAK/STAT, HIPPO and EGF/MAP^11–14^ pathway activity. Interestingly all of these proliferative pathways are dysregulated and contribute to ADPKD development^15–18^. Yet, the role of ANKHD1 ADPKD progression is unknown. To explore a potential role for ANKHD1 in the progression of ADPKD we examine its expression in human kidney sections from people with ADPKD and age and gender control healthy individuals.

Here we present the first mechanistic study identifying ANKHD1 as a master regulator of ADPKD disease progression. Specifically, to examine its potential involvement in human ADPKD *in vivo*, we performed immunostaining of human kidneys from patients with early to late stages of ADPKD and healthy control kidneys. To examine its role, we generated the first *Ankhd1*-deficient mice which we crossed with *Pkd1*^nl/nl^ mice to produce *Ankhd1*-deficient mice with ADPKD. To realize its mechanistic role in ADPKD, we performed transcriptome-wide profiling followed by next-generation RNA-sequencing of human patient derived cells with and without ANKHD1 siRNA. The data presented in this paper identify ANKHD1 as gene that promotes ADPKD disease progression.

## RESULTS

### ANKHD1 is expressed at all stages of human and murine ADPKD

Whether ANKHD1 is expressed in human ADPKD remains unknown. We performed immunohistochemical analysis of ANKHD1 expression and distribution in human kidney sections from patients with early to late stage ADPKD and compared this to kidneys derived from people without renal pathology (healthy control tissue). ANKHD1 was expressed highly at all stages of ADPKD (Fig 1a), whereas isotype control IgG antibody did not produce significant staining (fig 1c). To examine the level of ANKHD1 expression in disease versus healthy tissue, the immunostaining was quantified, and it was observed that ANKHD1 was highest at the earliest stages of disease but remained high through disease development (fig 1b). It is worthwhile to note that ANKHD1 expression was also identified in healthy non-ADPKD human kidney sections (fig 1a-b) but this expression was lower when compared with ADPKD (fig 1b). Hence, we conclude that ANKHD1 is temporally and spatially coincident with disease progression. To examine whether *Ankhd1* is also found at the site of disease in a mouse model of ADPKD, we performed immunohistochemistry of Ankhd1 in kidney sections of mice with Pkd1^nl/nl^ deficiency. Mouse Ankhd1 was highly expressed in Pkd1^nl/nl^ mice and its expression was associated with cyst lining epithelial cells (Fig 1d). Taken together this work identifies ANKHD1 as a protein that its expression spatially and temporally co-localised with ADPKD disease.

**Figure 1:**
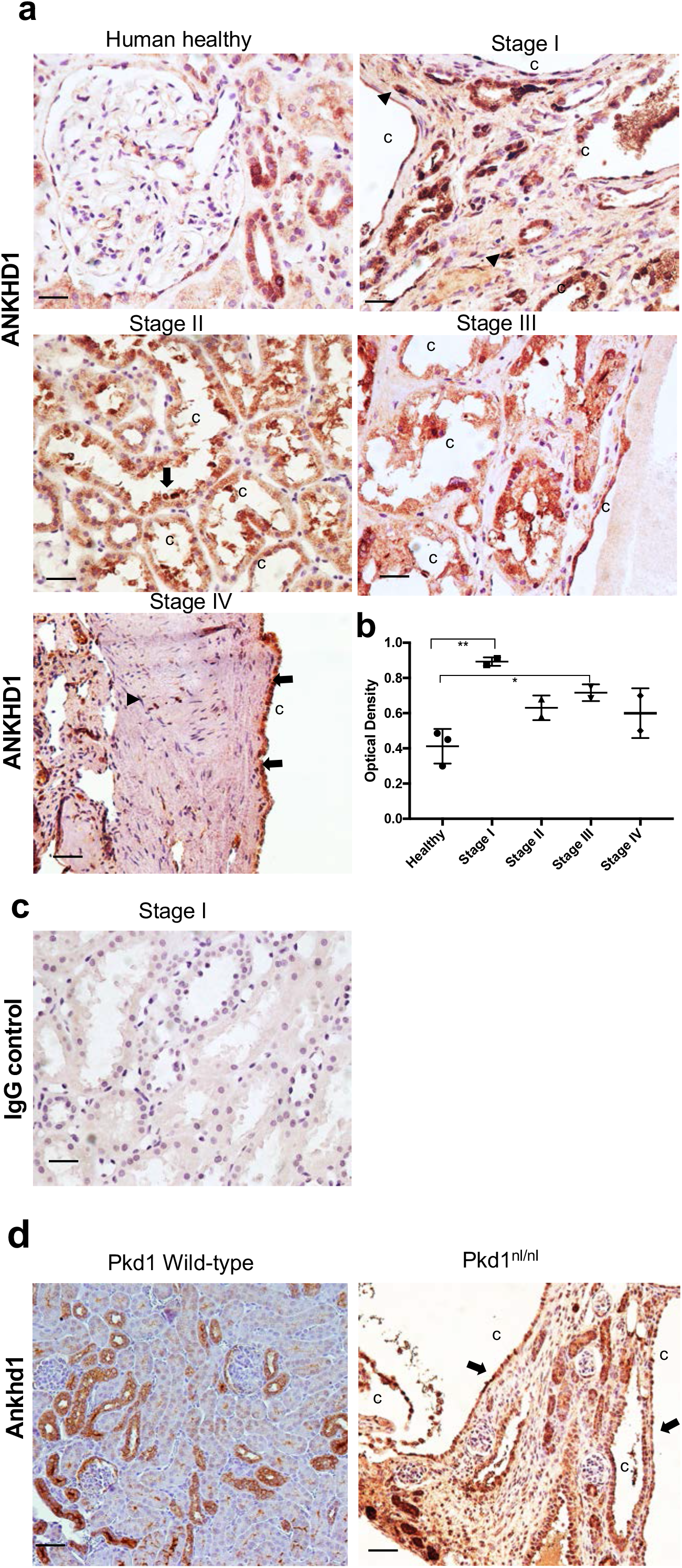
ANKHD1 is expressed at all stages of human and murine ADPKD. (a) DAB immunostaning of kidney sections from control healthy kidneys (n=3) and kidney from patients with different stages of ADPKD (n=8). Scale bar 100μm. (b) Graph shows quantification of the Ankhd1-DAB staining from healthy (n=3) and kidneys with stage 1 (n=2), stage II (n=2), stage III (n=2) and final stage IV (n=2) disease. Data are shown as ±s.e.m. One way ANOVA statistical analysis (**P<0.01, *P<0.05) with Bonferroni’s multiple comparison correction test. (c) Immunohistochemistry of Ankhd1 in kidney sections from control (left) and Pkd1nl/nl mice (4 weeks of age). Scale bar 50μm, c=cyst, arrow points at cyst lining epithelial cells and arrowheads at interstitial cells. (d) DAB immunostaining for ANKHD1 in wild type (left) and Pkd1nl/nl mice (right), with arrows pointing at ANKHD1 expression in cyst lining cells within cysts (c).

### *Ankhd1*-deficiency decelerates polycystic kidney disease development and fibrosis

To examine the potential involvement of ANKHD1 in the progression of ADPKD we used the first *Ankhd1*-deficient mice. Hemizygous *Ankhd1*^+/-^ mice were crossed with *Pkd1*^*nl*/+^ mice to generate *Ankhd1*^-/+^; *Pkd1^nl/nl^* (Ankhd1-deficient mice with polycystic kidney disease) and *Ankhd1*^+/+^; *Pkd1^nl/nl^* (Ankhd1 wild-type with polycystic kidney disease). We focused on mice with hemizygous deletion of the *Ankhd1* locus, since full *Ankhd1*-deficiency exhibited 100% embryonic lethality in the Pkd1 background, suggesting that complete knockout of *Ankhd1* is not compatible with life in mice with polycystic kidney disease. At 4 weeks of age, both *Ankhd1 Ankhd1*-deficient mice developed significantly reduced level of polycystic kidney disease in the Pkd1^nl/nl^ background (Fig 2a). Specifically, mice with *Ankhd1*-decifiency had significantly lower cystic index (fig 2b), their kidneys were lighter than wild-type littermate controls with disease (fig 2c) and developed fewer cysts overall (fig 2d), indicating that lowering the *Ankhd1* markedly reduced cystic disease burden. To examine whether lower cystic burden affected kidney function, blood urea nitrogen and kidney histology were analysed, and it was found that *Ankhd1*-deficiency led to a significant reduction in blood urea nitrogen (fig 2e) and improved renal histology compared to control animals with disease (fig 2f) signifying that *Ankhd1*-deficiency improves renal function in ADPKD. Another marker of disease progression in ADPKD is the development and progressive expansion of interstitial fibrosis (fibrosis around the tubular network, not associated with vessels). Given that ANKHD1 is involved in ADPKD progression, we measured picrosirius red staining to understand if it affects fibrogenesis in ADPKD. We found that *Ankhd1*-deficiency led to a significant reduction in serius red staining (fig 2g-h), suggesting that ANKHD1 inhibition may carry anti-fibrotic potential. Taken together the data from the first Ankhd1-deficient mice indicate that lowering ANKHD1 both blocks the extend of cystic burden as well as preserving renal function, compared to mice expressing wild-type levels of ANKHD1. These data provide the first evidence that Ankhd1-antagonism may provide a clinical benefit in ADPKD in two distinct ways, by slowing down cystic growth (and the damage associated with this, as per BUN) and by lowering the rate of fibrogenesis.

**Figure 2:**
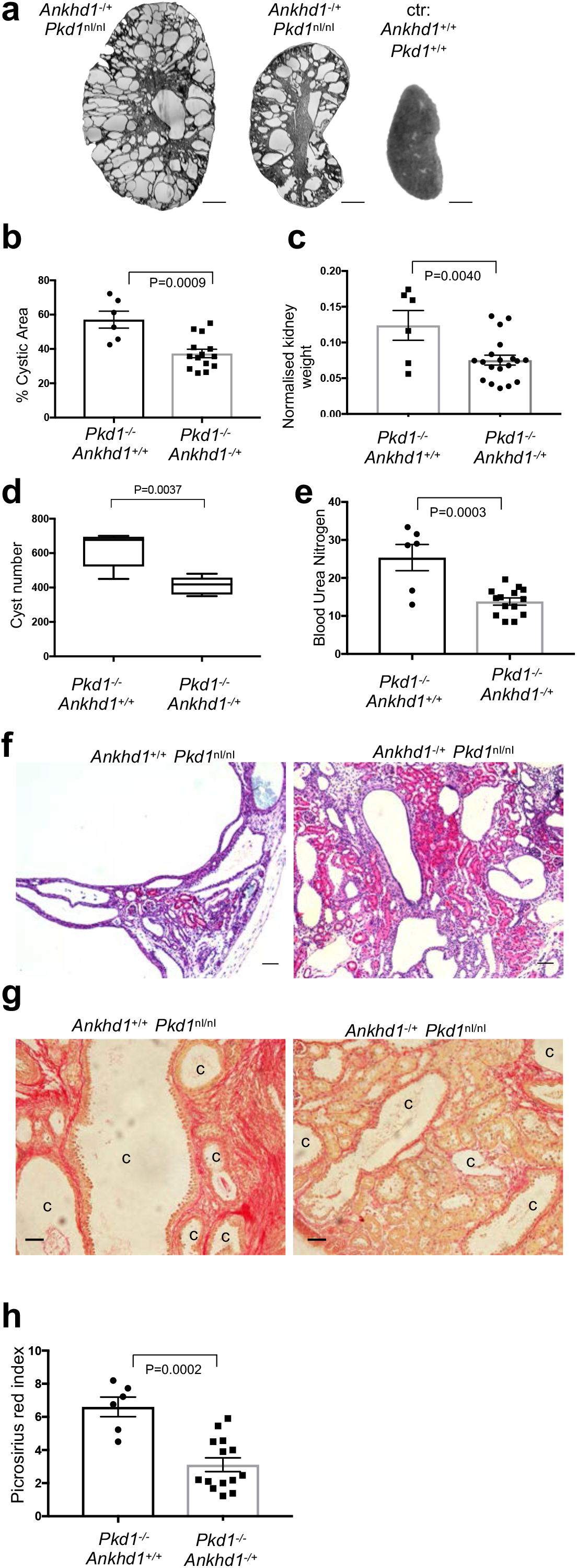
*Ankhd1*-deficiency decelerates polycystic kidney disease development and fibrosis. (a) Micrographs of H&E stained kidney sections from control mice with wild type *Pkd1* (*Pkd1*^+/+^) and *Pkd1*^nl/nl^ mice with hemizygous deletion of *Ankhd1*^-/+^ at 4 weeks of age. Scale bars at 1,000μm. The graphs show cystic index (b), the two kidney weight over body weight ratio (c), actual cyst number (d), and blood urea nitrogen (e) for controls and Ankhd1-deficient mice analysed at 4 weeks of age. Data are shown as ±s.e.m. T-test statistical analysis with indicated p values was performed. (f) H&E immunostaning was performed in control and Ankhd1-deficnet mice, scales are at 50μm, c=cyst. (g) Fibrosis was examined by picrosirius red staining and PSR index was quantified (h). Data are expressed as ±s.e.m and student t-tests were performed with indicated P values. Scales are at 100μm, c=cyst.

### Transcriptome-wide RNA-seq analysis identifies ANKHD1 as a major regulator of proliferation and extracellular matrix receptor interactions

To explore the mechanistic role of ANKHD1 in ADPKD, we performed next-generation RNA-sequencing in two human patient-derived renal epithelial lines in which we modulated the expression of ANKHD1 by RNA interference. Of the 11,614 genes detected, 534 genes showed statistically significant differential gene expression in cells with ANKHD1-siRNA (Ankhd1-low cells) when compared with cells transfected with control non-targeting siRNA (Ankhd1-high cells) (fig 3a), showing that the expression of these 534 genes is controlled by ANKHD1. We proposed that ANKHD1 may regulate important disease-controlling pathways relating to tissue growth or fibrosis, to identify these pathways we performed gene ontology analysis of the highly significant targets, identifying differentially expressed gene families involved in the control of tube formation and cell population proliferation (Fig 3b). Gene ontology clustering revealed a major role for ANKHD1 in the control of immune responses, especially via interferon and interferon signalling (Fig. 3c). To identify the functional pathways that are associated with *ANKHD1*-deficiency, we performed Kyoto Encyclopaedia of Genes and Genomes (KEGG) pathway enrichment analysis, which further revealed a role for ANKHD1 in the JAK/STAT signalling, cell-extracellular matrix (cell-ECM) interactions, TGFβ pathway and WNT signalling (Fig 3c). From the 534 genes, the expression of 21 genes with role in the regulation of JAK/STAT or cell-ECM interactions is shown (Fig 3c). Taken together, these results identify ANKHD1 as a master regulator of proliferation and matrix remodelling in ADPKD.

**Figure 3:**
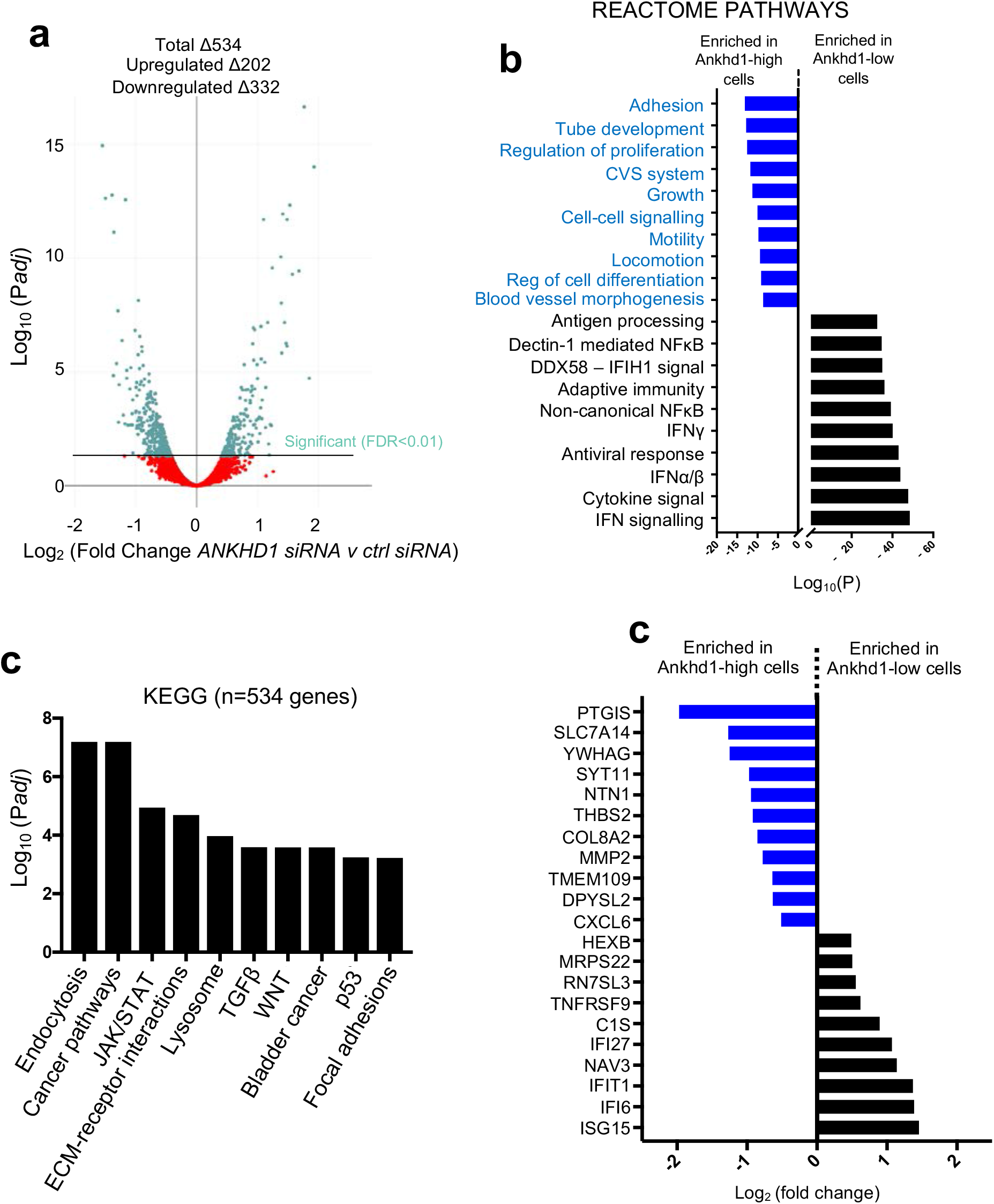
Transcriptome-wide RNA-seq analysis identifies ANKHD1 as a major regulator of proliferation and ECM-receptor interactions. (a) *ANKHD1*-siRNA whole transcriptome profiling in two biological samples (OX161c1 and SKI001) was performed using next-generation NextSeq500 Illumina platform. Volcano plots of genes with differential expression in Ankhd1-high (non-targeting siRNA) and Ankhd1-low cells (Ankhd1-siRNA) were generated (adjusted P ≤ 0.01 and fold change ≥ 0.5). P values were calculated with two-sided likelihood ratio test and adjusted by Benjamini-Hochberg method; n=4 (2 conditions Ankhd1-siRNA and nontarget siRNA and 2 biological samples). (b) Reactome pathway enrichment analysis of the n=534 differentially expressed genes in Ankhd1-high and Ankhd1-low cells identify Ankhd1 as a novel regulator of tube morphogenesis and cell proliferation in ADPKD cells. (c) KEGG analysis is shown for the 10 most statistically enriched pathways. (d) Expression of the top 21 genes involved in JAK/STAT or ECM remodelling is shown.

### ANKHD1 silencing reduces proliferation potential in human patient derived ADPKD cells

Since persistent proliferation is a hallmark of ADPKD progression yet the mechanisms underlying it are less clear, we set out to experimentally examine the potential role of ANKHD1 in the control of epithelial cell proliferation using immunostaining, flow-cytometry and cell cycle analysis in *Ankhd1*-deficient mice and in two human ADPKD lines in which we silenced human ANKHD1. Silencing of ANKHD1 caused more than 60% reduction in the number of Ki67 positive cells (fig 4a), suggesting that ANKHD1 is essential for proliferation of human ADPKD-derived epithelial cells. Likewise, the number of H3 (pser10) positive cells, representing cells undergoing mitosis, was significantly reduced in ANKHD1-siRNA cells (fig 4 b-d), this shows that ANKHD1 is an indispensable driver of mitosis in this disease context. To elucidate the underpinning mechanism, we performed cell cycle analysis, revealing that ANKHD1 silencing leads to a significant reduction of G2-M transitioning cells and is associated with an increase in G1 phase arrested cells (fig 4e), showing that ANKHD1 mechanistically controls proliferation by enhancing G2/M transitionThese data together identify ANKHD1 as an is indispensable gene for maintaining cell proliferation of ADPKD kidney epithelial cells.

**Figure 4:**
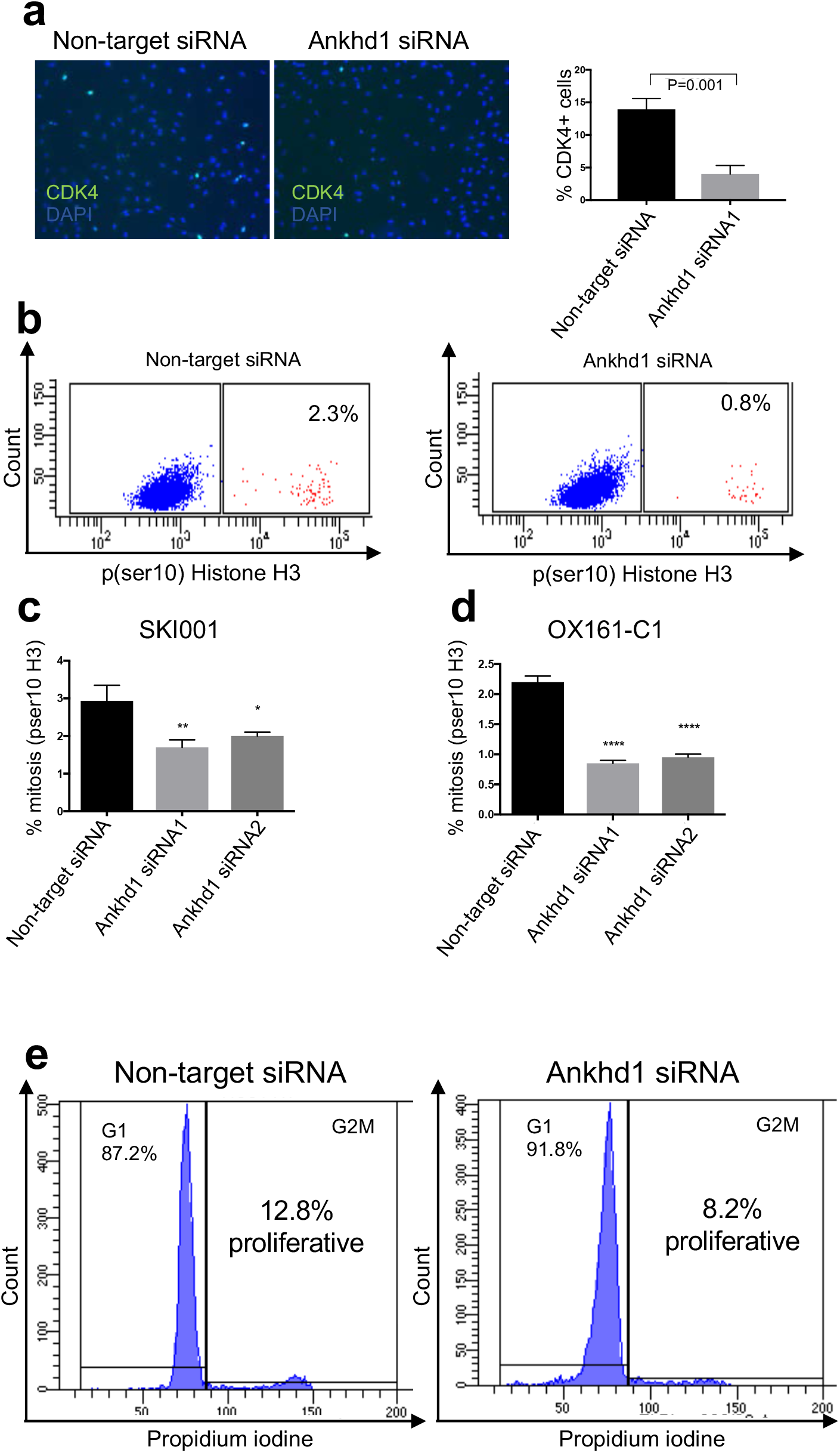
ANKHD1 silencing reduces proliferation in human patient-derived ADPKD cells. (a) SKI001 transfected with Ankhd1 siRNA show a decreased number of Ki67 positivity when compared with non-target siRNA transfected cells. Quantification of % of Ki67 positive cells is presented as ±s.e.m, student t-test was used to calculate the indicated p-value, n=3 biological replicates. (b) OX161-c1, transfected with Ankhd1 siRNA, exhibited decreased p(ser10) Histone H3 staining. (c-d) Quantification of the number of cells positive for p(ser10) histone H3 in SKI001 (c) and OXI161-c1 (d), data are shown as ±s.e.m, One way ANOVA statistical analysis (***P<0.001, **P<0.01, *P<0.05) with Bonferroni’s multiple comparison correction test. (e) Flow cytometry of SKI001 cells stained with propidium iodine was performed followed by cell cycle analysis.

### ANKHD1 enhances ADPKD proliferation by promoting STAT5 signalling

It was previously reported that ADPKD is characterised by altered JAK/STAT signalling and specifically STAT5 is elevated in polycystic kidneys^18^. Our previous RNA-seq data illustrated that ANKHD1 alters JAK/STAT activity, therefore, to investigate a potential role of Ankhd1 in the control of STAT5 signalling, we used mice with targeted *Ankhd1*-deficiency in the *Pkd1*^nl/nl^ model of polycystic kidney disease. To assess the role of ANKHD1 in the regulation of the STAT5 signalling pathway, we analysed phosphorylation, gene transcription and also performed immunostaining in kidneys of mice with ADPKD. Stat5 tyrosine phosphorylation was readily detected in *Pkd1*-deficient polycystic kidneys but was below detection level in wild-type uninjured kidneys. Intriguingly, Stat5 phosphorylation was significantly reduced in *Ankhd1*^-/+^ polycystic kidneys (fig 5a), when compared to *Ankhd1*^+/+^. Tyrosine phosphorylation of Stat5b (on Tyr 694) is required for the dimerization and transcriptional activity of this transcription factor^21^. Consistent with the diminished phosphorylation of Stat5b, its nuclear localisation was also reduced in *Ankhd1*-deficient polycystic kidneys (fig 5b). Furthermore, the induction of Insulin growth factor 1 (Igf1), CIS and Suppressor of cytokine signalling 3 (SOCS3), genes transcriptionally activated by Stat5^22^, were reduced in *Ankhd1*^-/+^ compared with *Ankhd1* control kidneys (Figure 5c). These results indicate that Ankhd1 positively regulates the STAT5 activity in ADPKD. Given that STAT5 is a core regulator of cell growth, we examined whether ANKHD1 can control proliferation of ADPKD cells. To examine whether Ankhd1 controls proliferation of cystic epithelial cells *in vivo*, we performed cyclin D1 immunostaining in kidney sections of *Ankhd1*^-/+^ and wild-type mice with polycystic kidney disease. Ankhd1-deficient mice showed significantly less cyclin D1 expression when compared with *Ankhd1*^+/+^ mice (fig 5d), illustrating that lowering Ankhd1 is sufficient to reduce proliferation of cystic cells *in vivo*. Collectively these data identify a novel role for *Ankhd1* in the regulation of the Stat5 proliferative signalling in the kidney.

**Figure 5:**
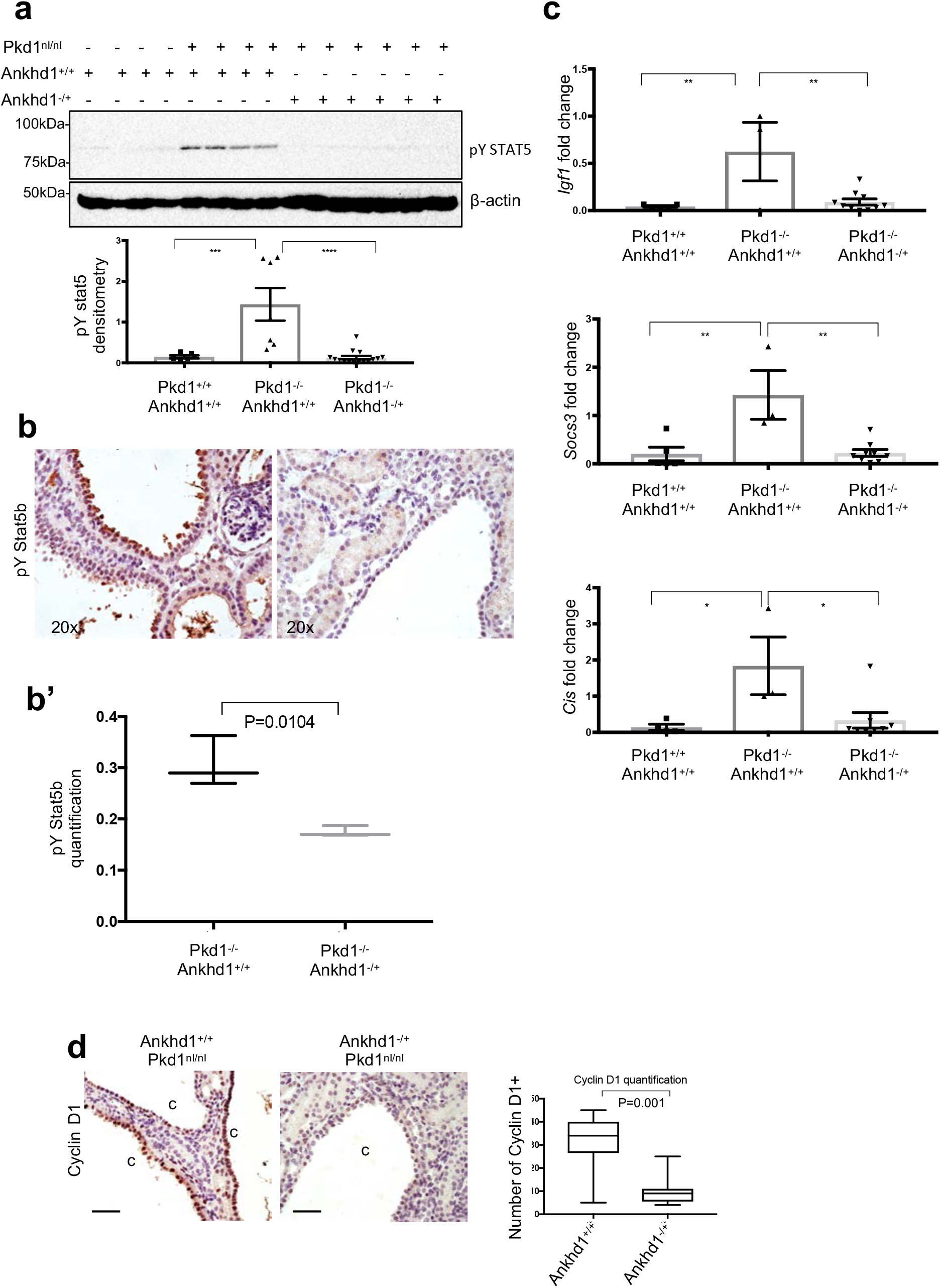
ANKHD1 enhances STAT5 transcriptional activity. (a) Decreased STAT5 tyrosine phosphorylation in mouse kidneys with Ankhd1-deficiency. Phosphorylation in *Ankhd1*^+/+^ and *Ankhd1*^-/+^ whole kidney lysates from *Pkd1^nl/nl^* with polycystic kidney disease and Pkd1 wild type (control) was determined with phosphospecific antibodies against Stat5 (Tyr 694) (pY Stat5). Quantification of pY Stat5 was performed by densitometry analysis and data are presented as as ±s.e.m. One-way ANOVA statistical analysis (***P<0.001, ****P<0.0001) with Bonferroni’s multiple comparison correction test was performed. (b) Decreased nuclear pY STAT5 was observed in Ankhd1-/+, quantification data (b’) as shown as as ±s.e.m and P value calculated using the student t–test. (c) Reduced induction of Igf1, Socd3 and Cis in mouse kidneys with hemizygous Ankhd1-deficiency. Induction of Igf1, Socs3 and Cis was observed in kidneys of mice with polycystic kidney disease (Pkd1^nl/nl^) by qPCR at 4-weeks of age, using Rpl13a as an internal control. Data as shown as as ±s.e.m. One-way ANOVA statistical analysis (**P<0.01, *P<0.05) with Bonferroni’s multiple comparison correction test was performed. (d) Cyclin D1 protein is decreased in mouse kidneys that lack Ankhd1 (Ankhd1-/+), determined by immunofluorescence staining using specific anti-cyclin D1 antibodies (brown), nuclear counterstain was achieved by haematoxylin staining (blue). Quantification of cyclin D1 was performed and data are presented as as ±s.e.m, student t-test was used to calculate the indicated p-value. Scale bars are 50μm, c=cyst.

## DISCUSSION

Here we describe the identification and characterisation of ANKHD1 as a master regulator of polycystic kidney disease progression and demonstrate that it plays key roles in proliferation and extracellular matrix remodelling. Proliferation and fibrosis are two critical processes contributing to ADPKD disease progression. Mechanistically we identify that ANKHD1 alters proliferative signalling by enhancing STAT5-driven transcription in ADPKD kidneys. This is interesting since it was recently shown that STAT5 is a major contributor to proliferation and ADPKD disease progression ^18^. The fact that ANKHD1 drives proliferation and enhances fibrogenesis together suggest that it is an ideal future therapeutic target, since its successful inhibition is predicted to block both the growth of cysts and inhibit tissue scarring, which is a final common pathway to organ failure irrespective of the initial cause of damage. This work therefore carries also significant implications for other disease characterised by scarring and proliferation.

The observation that *Ankhd1* is expressed during all stages of human ADPKD and in mouse models of the disease, indicates that *Ankhd1* is a key gene involved in cystic disease. Our findings are likely to be relevant to human ADPKD because *ANKHD1* inhibition also attenuated proliferation of human ADPKD renal cells and importantly ANKHD1 protein is highly expressed in human ADPKD biopsies at all stages of disease progression. Since ANKHD1 controls the ability of cells to divide, it is possible that its inhibition may slow the growth of cyst in additional ciliopathies ^23,24^ and other kidney diseases defined by epithelial overgrowth such as renal cancer^25^, besides ADPKD.

Mechanistically at present, the insights into *Ankhd1-induced* pathways mainly focus on its effects in controlling the HIPPO organ-size control pathway^26–28^ and the EGF/MAP kinase pathway ^12–14^ However, some investigations have suggested roles for the *Drosophila Ankhd1* in JAK/STAT signalling ^11^. Our current molecular and transcriptome-wide observations are consistent with a potential role for mammalian *Ankhd1* in the control of the JAK/STAT pathway. Hence, these insights further our understanding of the underlying control mechanisms of a core signalling pathway and indicate that mammalian ANKHD1 is a novel regulator of STAT5.

ANKHD1-deficiency led to a significant reduction in fibrosis in the polycystic kidney. Since fibrosis is a major problem underlying many pathologies of various organs (liver, lung, skin etc), our finding placing ANKHD1 as a regulator of ECM remodeling may be of benefit to additional pathologies. Next-generation RNA-sequencing data identified a distinct ANKHD1-driven ECM remodelling signature that is altered when ANKHD1 is silenced, thus supporting a role for ANKHD1 in the control of fibrotic processes. It has not escaped our attention that ANKHD1-deficiency leads to enhanced interferon responses, with interferon being the most significant functional hit for ANKHD1, presented here. This is intriguing since strong evidence exists suggesting that interferon alpha therapy suppresses fibrosis^29^, hence it is hypothesized that the fibrotic effects of ANKHD1 may be mediated by IFN inhibition. While beyond the remit of this study, an in-depth analysis of the interplay of ANKHD1 and interferon pathway is warranted.

The results of this study provide the first genetic, functional and biochemical evidence that ANKHD1 promotes polycystic kidney disease progression and fibrosis by directly controlling epithelial cell proliferation, cytokine receptor signalling and ECM remodelling. Genetic inactivation of the *ANKHD1* gene in human cells and in mouse kidneys, inhibited proliferation and cytokine-triggered STAT5 transcriptional activities and blocked G2-M transitioning epithelial cells. These data identify ANKHD1 as an indispensable positive regulator of epithelial cells proliferation, fibrosis and cytokine receptor signalling highly relevant to polycystic kidney disease and further demonstrate a potential utility to other diseases characterised by abnormal proliferation and/or JAK/STAT signalling.

## Methods

### Mice

Previously described Pkd1^nl/nl^ mice ^30^ were bred with Ankhd1^-/+^ for this study under the 70/8968 UK home office project licence. For all studies an equal number of males and females were used. The mice listed above were maintained in the C57Bl6 background and their ages are indicated in the results section or figure legends.

### Human specimens

4% paraformaldehyde-fixed paraffin-embedded slices from normal and ADPKD kidneys were provided by the UK PKD Charity – sponsored Biobank. The fully-anonymised human samples were procured at source in the OR after sterile perfusion immediately post-mortem or at time of nephrectomy by National Institute’s of Health (NIH) ethically-approved National Disease Research Interchange (NDRI, Philadelphia, USA) with full Institutional Review Board (IRB)/NIH approval in USA and in UK under the auspices of UCL ethical approval (Number 05/Q0508/6).

### RNA isolation and qPCR

Total RNA was isolated from cultured cells or mouse kidneys using the total RNA extraction miRNeasy mini kits (Qiagen). cDNA synthesis was performed using the iScript cDNa synthesis kit (Bio-rad), and qPCR was performed using the iQ SYBR green supermix (bio-rad).

### RNA sequencing

SKI001 and OX161-c1 human ADPKD-derived cells were treated with either siRNA against ANKHD1 (pool of 4 siRNAs) or a non-targeting siRNA (2 cell types x 2 treatments). RNA was extracted was subjected to quality checks, after which libraries were sequenced at the Cambridge Genomics Services (CGS) facility based within the University of Cambridge (UK), using a NextSeq500 Illumina sequencer to generate 75bp paired-end reads. Quality control of the reads was done using FastQC (v0.11.4). Reads were trimmed to remove low-quality bases before mapping, using TrimGalore (v0.4.1). Trimmed reads were mapped to the human genome with STAR (v2.5.2a), using the annotated transcripts from the ensembl GRCh38.95 GTF. The number of reads that mapped to genomic features was calculated using HTSeq-count (v0.6.0). Differential gene expression analysis was done using edgeR (3.26.5). Genes with an FDR corrected p-value of either <0.01 (siRNA experiment) or <0.05 (RIP-seq) were deemed significantly differentially expressed between the specified conditions. The identified genes were analysed using Gene Set Enrichment analysis for hallmark pathways or KEGG pathways (as specificed in figure legends) and molecular functions.

### Immunostaining

For immunostaining of 4& paraformaldehyde-fixed paraffin-embedded kidney sections or human cells grown on coverslips the following antibodies were used: cyclin D and KI67 antibodies were from Cell signalling (55506; 9449), stat5a, stat5b and Ankhd1 were from sigma (SAB2102320; SAB43000329; HPA008718). Quantification of staining was performed by randomly selecting 4 10x magnification fields per sample and subsequently determining positivity using Image J.

### Immunoblotting

Total protein was extracted from cells and kidneys and 20μg were loaded onto a 4-20% gradient SDS-polyacrylamide gel and proteins transferred onto a nitrocellulose membrane. PYstat5 and b-actin antibodies were from Cell signalling (9359 and 3700). Secondary antibodies anti-mouse-HRP and anti-rabbit HRP were from Dako and membranes were developed using ECL prime and imaged in Bio-Rad imager using software from Bio-Rad.

### Flow cytometry

Flow cytometry was used to assess cell cycle using propidium iodine on methanol fixed cells and mitosis using a phosphorylated serine 10 histone H3 from cell signalling (8552s).

### Statistical analysis

Data are shown as the mean +/-S.E.M. Statistical analysis was performed using either the student-t-test or one-way Anova followed by Tukey’s post hoc test to correct for multiple testing. P values < 0.05 were considered significant.

The authors declare that they have no conflict of interest

